# Longitudinal immunosequencing in healthy people reveals persistent T cell receptors rich in public receptors

**DOI:** 10.1101/262667

**Authors:** Nathaniel D. Chu, Haixin Sarah Bi, Ryan O. Emerson, Anna M. Sherwood, Michael E. Birnbaum, Harlan S. Robins, Eric J. Alm

**Author notes:** Corresponding author: Email addresses: NDC, HSB, ROE, AMS, MB, HSR, EJA.

## Abstract

**Background:** The adaptive immune system maintains a diversity of T cells capable of recognizing a broad array of antigens. Each T cell’s specificity and affinity for antigens is determined by its T cell receptors (TCRs), which together across all T cells form a repertoire of tens of millions of unique receptors in each individual. Although many studies have examined how TCR repertoires change in response to disease or drugs, few have explored the temporal dynamics of the TCR repertoire in healthy individuals.

**Results:** Here we report immunosequencing of TCR β chains (TCRβ) from the blood of three healthy individuals at eight time points over one year. TCRβ repertoires from samples of all T cells and memory T cells clearly clustered by individual, confirming that TCRβ repertoires are specific to individuals across time. This individuality was absent from TCRβs from naive T cells, suggesting that these differences result from an individual’s antigen exposure history. Many characteristics of the TCRβ repertoire (e.g., alpha diversity, clonality) were stable across time, although we found evidence of T cell expansion dynamics even within healthy individuals. We further identified a subset of “persistent” TCRβs present across all time points, and these receptors were rich in clonal and public receptors.

**Conclusions:** Our results revealed persistent receptors that may play a key role in immune system maintenance. They further highlight the importance of longitudinal sampling of the immune system and provide a much-needed baseline for TCRβ dynamics in healthy individuals. Such a baseline should help improve interpretation of changes in the TCRβ repertoire during disease or treatment.

## BACKGROUND

T cells play a vital role in cell-mediated immunity, one branch of the adaptive immune response against foreign and self-antigens. Upon recognizing an antigen from an antigen-presenting cell, naive T cells activate and proliferate rapidly. This process stimulates an effector response to the immediate challenge, followed by generation of memory T cells, which form a lasting cohort capable of mounting more-efficient responses against subsequent challenges by the same antigen.

The key to the flexibility and specificity of T cell responses lies in the cells’ remarkable capacity to diversify their T cell receptor (TCR) sequences, which determine the antigens those cells will recognize. Most T cells display TCRs made up of two chains: an a and a b chain. Sequence diversity in these chains arises during T cell development, through recombination of three sets of gene segments: the variable (V), diversity (D), and joining (J) segments (1). Random insertions and deletions at each genetic junction introduce still more diversity, resulting in a theoretical repertoire of 10^15^ unique receptors in humans (2). Selective pressures during and after T cell development, as well as constraints on the number of T cells maintained by the body, limit this diversity to an observed 10^7^ (approximately) unique receptors per individual (2–5).

This TCR repertoire forms the foundation of the adaptive immune response, which dynamically responds to disease. Each immune challenge prompts expansions and contractions of different T cell populations, and new T cells are continually generated. Substantial research interest has focused on these dynamics in the context of immune system perturbations, including in cancer (6–9), infection (10,11), autoimmune disorders (12,13), and therapeutic trials (8,14,15). Observing changes in TCR populations not only uncovers cellular mechanisms driving disease, but can inform development of new diagnostics, biomarkers, and therapeutics involving T cells.

Less research has explored TCR dynamics in healthy individuals. Previous studies found that some TCRs remain present in individuals over decades (16,17), but these long-term studies may not directly relate to shorter-term events, such as diseases or treatments. Interpreting TCR dynamics when the immune system is challenged would be more straightforward if we had a clear picture of TCR dynamics in healthy individuals.

To help develop this picture, we report immunosequencing of peripheral TCR β chain (TCRβ) repertoires of three individuals at eight time points over one year. We focused on the TCRβ chain because, unlike the α chain, only one β chain can be expressed on each T cell (18), the β chain contains greater sequence diversity (19), and it more frequently interacts with presented antigens during recognition (20). These factors suggest that TCRβ sequences should be sufficient to track individual T cells and their clones. Our analysis revealed overall individuality and temporal stability of the TCRβ pool. We also uncovered a set of temporally persistent TCRβs, which were more abundant, and shared across more people, than transitory TCRβs.

## RESULTS

### T cell receptor repertoires show individuality and stability through time

To characterize the dynamics of T cell receptors in healthy individuals, we deeply sequenced the TCRβ locus of peripheral blood mononuclear cells (PBMCs) isolated from three healthy adults (for schematic of experimental design, see **Figure 1a**). We sampled each individual at eight time points over one year (**Figure 1a**). For three intermediate time points, we also sequenced flow-sorted naive and memory T cells from PBMCs (see Methods). We summarize per-sample sequencing reads, unique TCRβs—which we defined as a unique combination of a V segment, CDR3 amino acid sequence, and J segment (21)—and other global statistics in **Table S1**. Most TCRβs had abundances near 10^-6^ (**Figure S1**), and rarefaction curves indicate that all samples were well saturated (**Figure S2**). This saturation indicates that our sequencing captured the full diversity of TCRβs in our samples, although our blood samples cannot capture the full diversity of the TCRβ repertoire (see **Discussion**).

**Figure 1.**
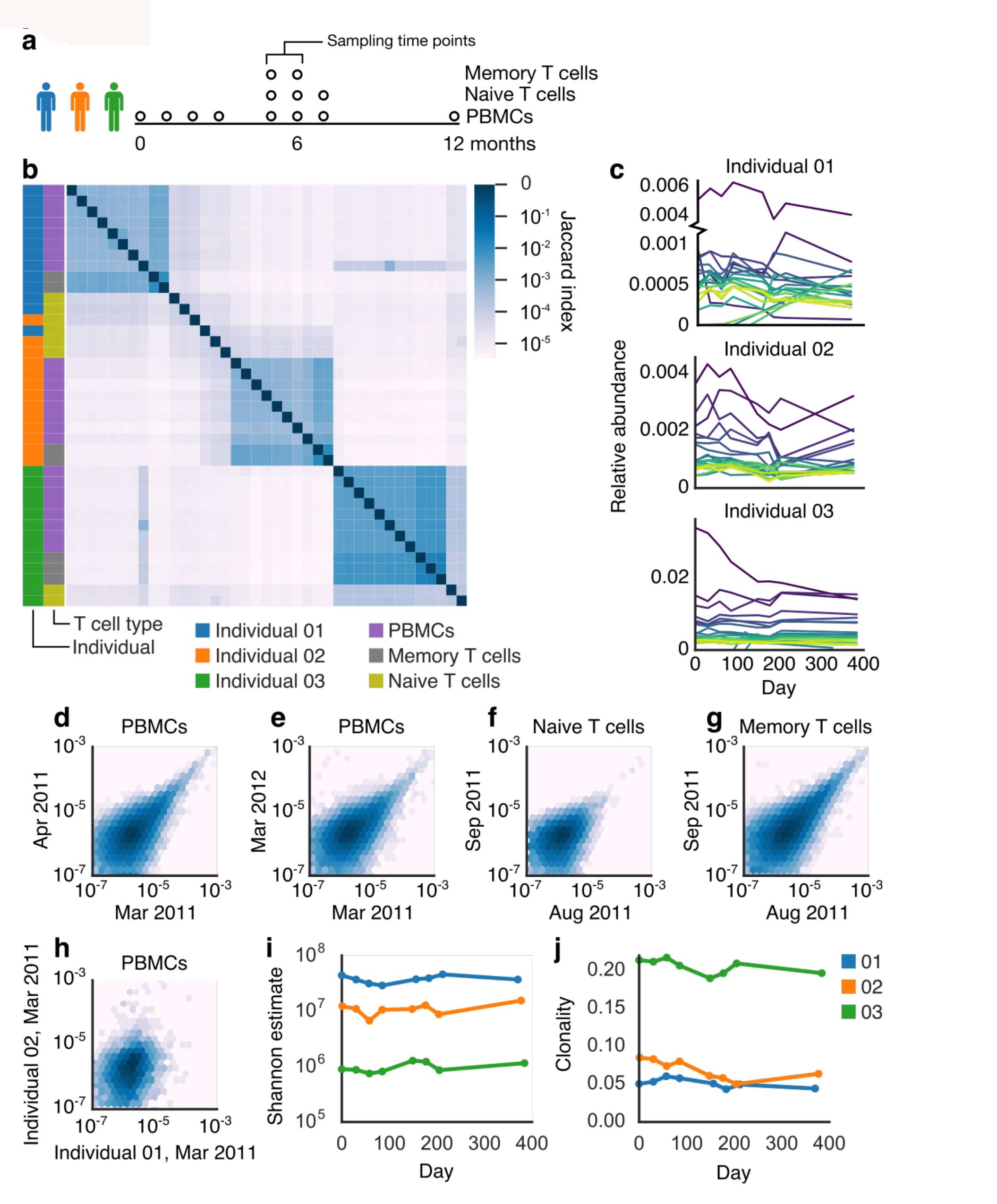
The TCRβ repertoire displayed stability and individual-specific characteristics. (**a**) Experimental design of T cell sampling. (**b**) A heatmap of Jaccard indexes shows clear clustering of samples by individual. Samples of naive T cells clustered less by individual than did PBMC or memory T cell samples. Relative abundances of the 20 most abundant TCRβs (**c**) appeared stable through time. TCRβ abundances in PBMCs correlated within an individual across time points, including across a month (**d**, shared TCRβs = 33601, Spearman *rho* = 0.55718,*p* < 10^−6^), and a year (**e**, shared TCRβs = 25933, Spearman *rho* = 0.53810, *p* < 10^−6^), as well as across a month in naive (**f**, shared TCRβs = 15873, Spearman *rho* = 0.37892, *p* < 10^−6^) and memory T cells (**g**, shared TCRβs = 47866, Spearman *rho* = 0.64934, *p* < 10^−6^). TCRβs correlated much less across individuals (**h**, shared TCRβs = 5014, Spearman *rho* = 0.28554, *p* < 10^−6^). Shannon alpha diversity estimate (**i**) and clonality (defined as 1 – Pielou’s evenness, **j**) of the TCRβ repertoire were consistent over time.

We first examined whether previously observed differences among individuals were stable through time (7,22). Looking at shared TCRβs (Jaccard index) among samples, we indeed found that samples of PBMCs or memory T cells taken from the same individual shared more TCRβs than samples taken from different individuals (**Figure 1b**), and this pattern was consistent over one year. In adults, memory T cells are thought to make up 60–90% of circulating T cells (23,24), which aligns with the agreement between these two T cell sample types. In contrast, TCRβs from naive T cells did not cluster cohesively by individual (**Figure 1b**). As naive T cells have not yet recognized a corresponding antigen, this lack of cohesion suggests that before antigen recognition and proliferation, TCRβ repertoires are not specific to individuals (**Figure 1b**). We can thus conclude that individuality results from an individual’s unique antigen exposure and T cell activation history.

We next examined patterns across samples from the same individual to understand TCR dynamics in healthy individuals. We observed only a minority of TCRβs shared among samples from month to month; indeed, samples of PBMCs at different months from the same individual typically shared only 11% of TCRβs (standard deviation 3.6%, range 5–18%) (**Figure 1b**).

Two factors likely played a role in the observed turnover of TCRβ repertoires: (1) changes in TCRβ abundances across time and (2) inherent undersampling of such a diverse system (see **Discussion**). Undersampling likely explained much of the low overlap of TCRβs among samples. To verify that patterns we observed were not artifacts of undersampling, we also analyzed a subset of high-abundance TCRβs (see **Methods**), which are less likely to be affected. In these TCRβs, we observed typical sharing of 63% (standard deviation 13.8%, range 35–88%) of TCRβs in PBMC samples across time (**Figure S3a**). PBMC and memory T cell samples (but not naive T cell samples) still clearly clustered by individual when only these TCRβs were considered (**Figure S3a**).

The frequencies of high-abundance TCRβs from each individual were largely consistent over time (**Figure 1c**). We found that abundances of the same TCRβs correlated within individuals over the span of a month (**Figure 1d, S3b**) and a year (**Figure 1e, S3c**). This correlation was particularly strong for more abundant TCRβs (**Figure S3b–c**) whereas rare TCRβs varied more. This correlation held true in naive and memory T cell subpopulations, sampled across a month (**Figure 1f-g**). In contrast, correlation was much weaker among abundances of TCRβs shared across individuals (**Figure 1h, S3d**), again highlighting the individuality of each repertoire. We found that the proportion of shared TCRβs (Jaccard index) tended to decrease with longer time intervals passed between samples, although with a notable reversion in Individual 02 (**Figure S4**). We observed stable alpha diversity (**Figure 1i, S3e**), clonality (**Figure 1j, S3f**), and V and J usage (**Figure S5, S6**) within individuals over time.

In the absence of experimental intervention, we observed complex clonal dynamics in many TCRβs, including cohorts of TCRβs with closely correlated expansion patterns (**Figure S7**). To avoid artifacts from undersampling, we looked for such cohorts of correlating receptors only in high-abundance TCRβs (see **Methods**). In all individuals, many high-abundance TCRβs appeared together only at a single time point. We also found cohorts of high-abundance TCRβs that correlated across time points (**Figure S7**). Some of these cohorts included TCRβs that fell across a range of abundances (**Figure S7a-b**), while other cohorts were made up of TCRβs with nearly identical abundances (**Figure S7c**). Correlating TCRβs were not obviously sequencing artifacts (**Table S2**, **Methods**). These cohorts of closely correlated TCRβs indicate that even in healthy individuals whose overall TCR repertoire appears stable, there remain underlying dynamics.

Taken together, these results revealed a diverse system, which nevertheless displayed consistent, unifying features differentiating individuals, plus longitudinal dynamics that suggested continual immune processes.

### A persistent TCRβ repertoire contains elevated proportions of clonal, public TCRβs

During our analysis, we discovered a subset of TCRβs was present across all PBMC samples from a single individual, a subset we called “persistent” TCRβs (**Figure 2a**). While approximately 90% of unique TCRβs in an individual’s PBMC sample from any given time point occurred only in that sample, 0.3–0.8% of TCRβs occurred at all eight time points (**Figure 2a**). When we considered only high-abundance TCRβs, up to 61% of high-abundance TCRβs appeared in only a single sample, while up to 88% appeared in all samples (**Figure S8a**).

**Figure 2.**
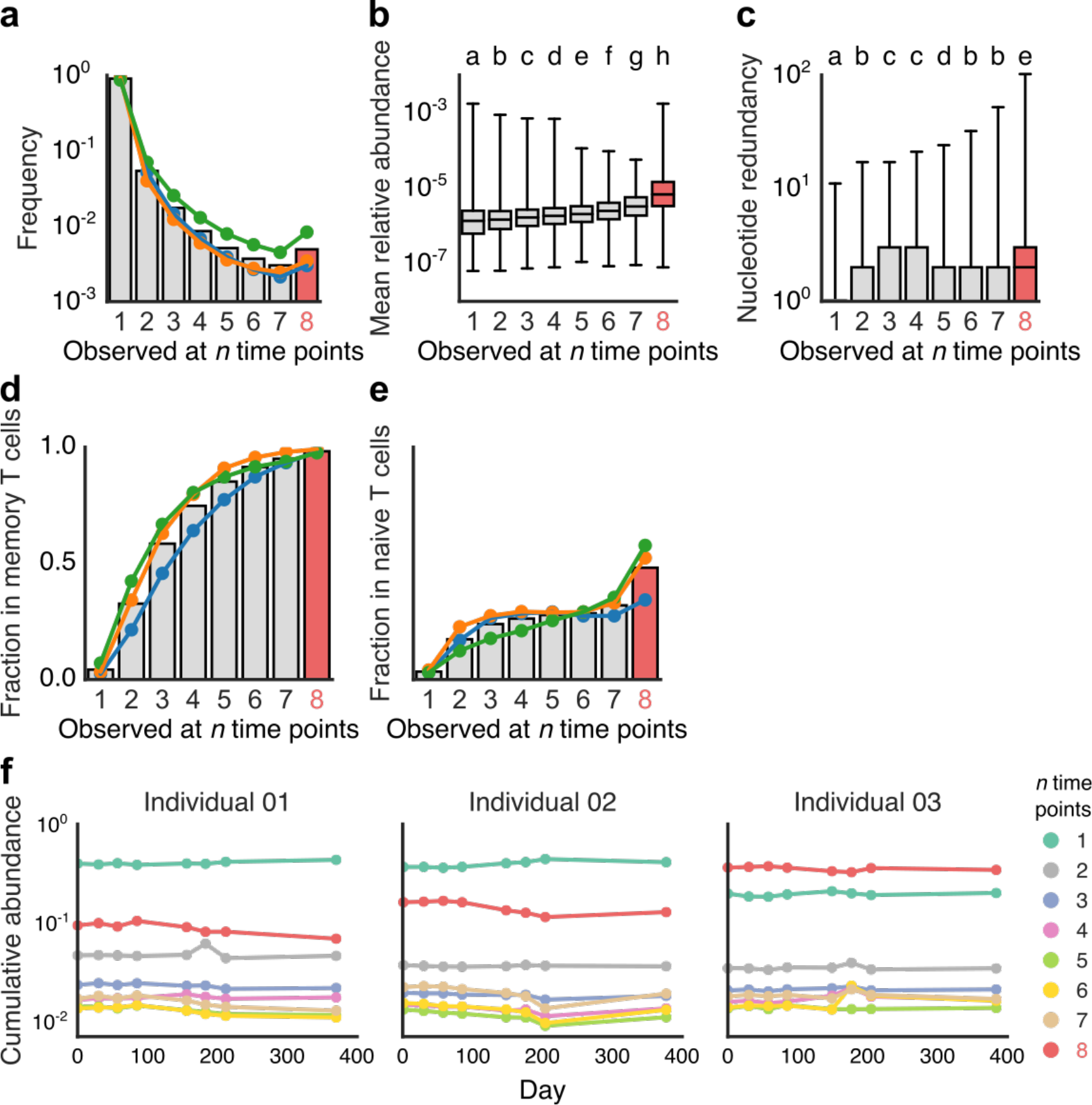
A subset of the TCRβ repertoire occurred across all time points—the persistent TCRβ repertoire. (**a**) The number of TCRβs observed at *n* time points. Persistent TCRβs tended to have greater abundance (**b**) and nucleotide sequence redundancy (**c**). Letters indicate significant differences by a Mann-Whitney *U* test (*p* < 0.001). Persistent TCRβs had higher proportions of TCRβs in common with memory (**d**) and with naive (**e**) T cell populations and constituted a stable and significant fraction of overall TCRβ abundance across time (**f**).

**Figure 3.**
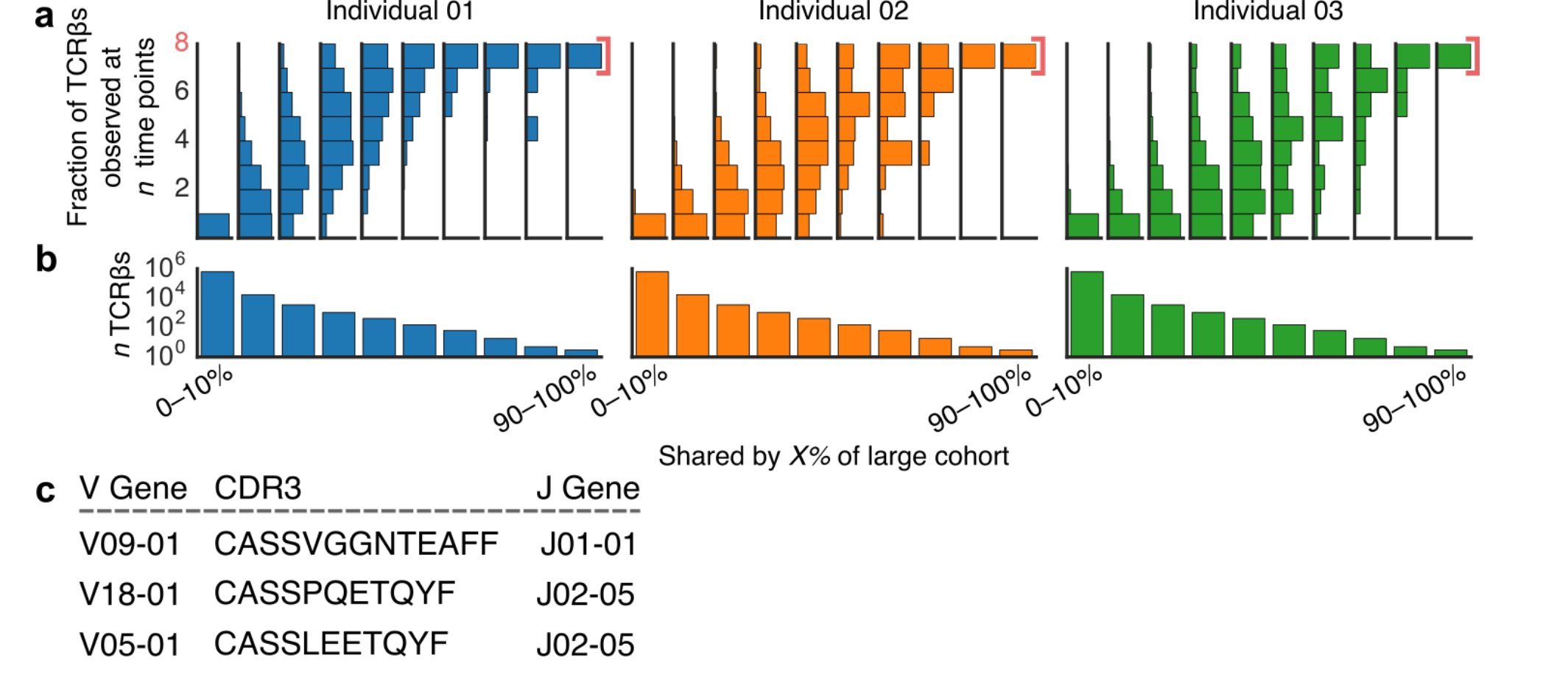
Public TCRβs are enriched in persistent TCRβs. We identified public TCRβs occurring in 0–10%, 0–20%, … 90–100% of individuals in a large cohort of similarly profiled subjects (*N* = 778). For each of these decile bins, we examined TCRβs shared with each of our three individuals’ time series data and tallied the number of time points at which we observed each TCRβ. (**b**) Vertical histograms of these distributions indicate that more private TCRβs—TCRβs shared by few people—most often occurred in only a single time point, while more public TCRβs tended to persist across time. (**c**) The number of TCRβs evaluated in each decile bin. (**d**) The three most public TCRβs (in over 90% of 778 individuals) were persistent in all three individuals.

We hypothesized that these persistent TCRβs might be selected for and maintained by the immune system, perhaps to respond to continual antigen exposures or other chronic immunological needs.

In our data, we found multiple signatures of immunological selection acting on persistent TCRβs. The members of this persistent subset tended to have a higher mean abundance than TCRβs observed at fewer time points (**Figure 2b**). We also observed that the number of unique nucleotide sequences encoding each TCRβ’s CDR3 amino acid sequence was generally higher for persistent TCRβs (**Figure 2c**). This pattern of greater nucleotide redundancy varied across individuals and region of the CDR3 sequence (**Figure S9a**), but TCRβs with the highest nucleotide redundancy were reliably persistent (**Figure S9b**). Furthermore, we discovered that TCRβs occurring at more time points, including persistent TCRβs, shared larger proportions of TCRβs also associated with memory T cells (**Figure 2d**). Remarkably, 98% of persistent TCRβs also occurred in memory T cells, suggesting that almost all persistent T cell clones had previously encountered and responded to their corresponding antigens. We found a similar pattern in naive T cells, although the overall overlap was lower (98% versus 50%) (**Figure 2e**). Persistent TCRβs did not show altered CDR3 lengths or VJ usage (**Figure S10–S12**). Like alpha diversity and clonality, the cumulative abundance of TCRβs present in different numbers of samples appeared stable over time and specific to individuals (**Figure 2f**). Surprisingly, although persistent TCRβs constituted less than 1% of all unique TCRβs, they accounted for 10–35% of the total abundance of TCRβs in any given sample (**Figure 2f**), further evidence that these T cell clones had expanded. We observed similar patterns when analyzing only high-abundance TCRβs (**Figure S8**).

Taken together, these characteristics—persistence across time, higher abundance, redundant nucleotide sequences, and overlap with memory T cells—suggest immunological selection for persistent TCRβs. We therefore investigated whether persistent TCRβs coexisted with TCRβs having very similar amino acid sequences. Previous studies have suggested that TCRβs with similar sequences likely respond to the same or similar antigens, and such coexistence may be evidence of immunological selection (25,26).

To explore this idea, we applied a network-based clustering algorithm based on Levenshtein edit distance between TCRβ CDR3 amino acid sequences in our data %(25–27). We represented antigen-specificity as a network graph of unique TCRβs, in which each edge connected a pair of TCRβs with putative shared specificity. We found that TCRβs having few edges—and thus few other TCRβs with putative shared antigen specificity—tended to occur in only one sample, while TCRβs with more edges included a higher frequency of TCRβs occurring in more than one sample (**Figure S13**, *p* < 10^−5^ for all three individuals by a nonparametric permutation test). This pattern indicates that TCRβs occurring with other, similar TCRβs were more often maintained across time in the peripheral immune system.

We next examined the association between persistent TCRβs—those shared across time points— and “public” TCRβs—those shared across people. Public TCRs show many of the same signatures of immunological selection as persistent TCRβs, including higher abundance (28), overlap with memory T cells (28), and coexistence with TCRs with similar sequence similarity (25). To identify public TCRβs, we compared our data with a similarly generated TCRβ dataset from a large cohort of 778 healthy individuals (21). We found that the most-shared (i.e., most-public) TCRβs from this large cohort had a larger proportion of persistent TCRβs from our three sampled individuals (**Figure 3a–b**, *p* < 10^−5^ for all three individuals by a nonparametric permutation test). Private TCRβs—those occurring in few individuals—most often occurred at only a single time point in our analyses. The three most public TCRβs (found in over 90% of the 778-individual cohort) were found to be in the persistent TCRβ repertoires of all three individuals and were diverse in structure (**Figure 3c**).

Public TCRs are thought to be products of genetic and biochemical biases in T cell receptor recombination (29,30) and also of convergent selection for TCRs that respond to frequently encountered antigens (21,32). To better understand the effects of biases during TCRβ recombination on receptor persistence, we used IGoR to estimate the probability that each TCRβ was generated before immune selection (33). Similar to previous studies (30), the probability that a given TCRβ was generated correlated closely with publicness (**Figure S14a**). In our time series data, TCRβs that occurred at multiple time points tended to have slightly higher generation probabilities (**Figure S14b**), but more-abundant TCRβs (both persistent and nonpersistent) did not (**Figure S14c–d**). These results suggest that, like public receptors, persistent receptors may partially result from biases in TCR recombination but that T cell abundance does not. Thus, although these two subsets of the TCR repertoire—persistent and public—are distinct, they overlap and share many characteristics, suggesting that both play a key role in immunity.

## DISCUSSION

Our analyses revealed both fluctuation and stability in the TCRβ repertoire of healthy individuals, providing a baseline framework for interpreting changes in the TCR repertoire. We identified a number of consistent patterns (e.g., alpha diversity, clonality), which are known to be affected by immunizations, clinical interventions, and changes in health status (7,14,34). These patterns differed among individuals across time, highlighting the role played by genetics and history of antigen exposure in shaping the TCR repertoire.

We further discovered a subset of persistent TCRβs that bore signs of immune selection. Persistent TCRβs tended to be more abundant than nonpersistent receptors, although this distinction is to a certain extent confounded by the fact that high abundance receptors are also more likely to be detected in a given sample. Nevertheless, this circular logic does not detract from the immune system’s maintenance of specific dominant TCRβs across time. We further found that persistent TCRβs had higher numbers of distinct nucleotide sequences encoding each TCRβ. TCR diversity is generated by somatic DNA recombination, so it is possible for the same TCR amino acid sequence to be generated from independent recombinations in different T cell clonal lineages. Thus, coexistence of multiple clonal lineages encoding the same TCRβ amino acid sequence may reflect selective pressures to maintain that TCRβ and its antigen specificity. Similarly, the presence of many TCRβs similar to persistent TCRβs—as identified by our network analysis—could also result from selection for receptors that recognize a set of related antigens (20,35). Previous studies using network analyses also found that public TCRβs tend to occur with similar TCRβs (25), further suggesting that both public and persistent TCRβs are key drivers of lasting immunity. In addition to using TCRβ sequencing to track TCRβs that proliferate in response to intervention, we propose that these two dimensions—publicness across individuals and persistence through time—represent two useful strategies for identifying biologically important TCRβs.

The presence of very public (present in >90% of individuals in our cohort) and persistent TCRβs led us to speculate that these TCRβs might be responding to a set of common antigens repeatedly encountered by healthy people. These antigens could be associated with self-antigens, chronic infections (e.g., Epstein-Barr virus), or possibly members of the human microbiota. In fact, CDR3 sequence CASSPQETQYF has been previously associated with the inflammatory skin disease psoriasis (36) and CASSLEETQYF has been implicated in responses to *Mycobacterium tuberculosis* (20) and cytomegalovirus (37).

In addition to persistent TCRβs, our analysis revealed many receptors with unstable behavior. Many high-abundance TCRβs did not persist through time, with many occurring at only a single time point (**Figure 2b, S8a**). These TCRβs could well correspond to T cells that expanded during a temporary immune challenge but then did not persist in high abundance afterward. The presence of dynamically expanding TCRβs in apparently healthy individuals poses an important consideration for designing studies monitoring the immune system. Studies tracking TCR abundances in cross-sectional immune system sampling (7,14,34–36) may capture not only T cell clones responding to intervention, but also expanding clones inherent in the T cell dynamics of healthy individuals. Repeated sampling before and after intervention could minimize such false positives.

Current immunosequencing methods have limitations that should inform the interpretation of our results. Most important, given such a diverse system as the TCR repertoire, even large sequencing efforts like ours undersample. Although our sequencing appeared to saturate our samples (**Figure S2**), additional bottlenecks during library preparation and, particularly, blood sampling limit our ability to capture full TCRβ diversity. Previous studies exhaustively sequenced multiple libraries from multiple blood samples, but even these estimates are considered a lower limit of TCRβ diversity (41). This detection limit could confound our identification of persistent TCRβs. Many of the TCRβs that did not occur in all samples were undoubtedly present but too rare for our analysis to capture. Thus, identification of a persistent TCR repertoire was subject to an abundance cutoff, whereby we focused on TCRs that persisted above the detection limit of sampling. To check that our conclusions were not heavily altered by undersampling, we analyzed high-abundance TCRβs and found similar overall patterns, so we infer that our main conclusions are likely robust despite this experimental limitation.

## CONCLUSIONS

To better understand healthy immune system dynamics in humans, we profiled the TCRβ repertoires from three individuals over one year. We found a system characterized by both fluctuation and stability and further discovered a novel subset of the TCRβ repertoire that might play a key role in immunity. As immune profiling in clinical trials becomes more prevalent, we hope that our results will provide much-needed context for interpreting immunosequencing data, as well as for informing future trial designs.

## METHODS

### Sample collection

Three healthy adult female volunteers ages 18–45 provided blood samples over the course of one year, with samples taken on a starting date and 1, 2, 3, 5, 6, 7, and 12 months after that date (**Figure 1a**). We sequenced TCRβ chains from approximately 1 million PBMCs from each sample. From the samples at 5, 6, and 7 months, we also sequenced TCRβ chains from sorted naive (CD3+, CD45RA+) and memory (CD3+, CD45RO+) T cells.

### High-throughput TCRβ sequencing

We extracted genomic DNA from cell samples using a Qiagen DNeasy blood extraction kit (Qiagen, Gaithersburg, MD, USA). We sequenced CDR3 regions of rearranged TCRβ genes and defined these regions according to the international immunogenetics information system (IMGT) (42). We amplified and sequenced TCRβ CDR3 regions using previously described protocols (2,43). Briefly, we applied a multiplexed PCR method, using a mixture of 60 forward primers specific to TCR Vβ gene segments plus 13 reverse primers specific to TCR Jβ gene segments.

We sequenced 87 base-pair reads on an Illumina HiSeq System and processed raw sequence data to remove errors in the primary sequence of each read. To collapse the TCRβ data into unique sequences, we used a nearest-neighbor algorithm—merging closely related sequences—which removed PCR and sequencing errors.

### Data analysis

In our analyses, we focused on TCRβs containing no stop codons and mapping successfully to a V gene and J gene (**Table S1**). Relative abundances of these “productive” TCRβ sequences, however, took into account the abundances of nonproductive TCRβ sequences, as these sequences were still part of the greater TCRβ pool. We defined a TCRβ as a unique combination of V gene, J gene, and CDR3 amino acid sequence. We examined nucleotide redundancy of each TCRβ by counting the number of T cell clones—a unique combination of V gene, J gene, and CDR3 nucleotide sequence—encoding each TCRβ. We considered TCRβs whose abundances ranked in the top 1% for each sample as high-abundance TCRβs, and we analyzed these TCRβs in parallel with the full TCRβ repertoire as a check for artifacts of undersampling (**Figure S5, S8**).

We calculated Spearman’s and Pearson’s correlation coefficients for TCRβ abundances across samples using the Python package SciPy, considering only TCRβs that were shared among samples. We calculated alpha diversity (Shannon estimate = e^(Shannon entropy)^) and clonality (1 – Pielou’s evenness) using the Python package Scikit-bio 0.5.1. We calculated Levenshtein distance using the Python package Python-Levenshtein 0.12.0 and analyzed the resulting network using the Python package NetworkX 1.9.1.

To look for TCRβs with similar temporal dynamics, we focused on TCRβs that occurred in high-abundance at least twice. These TCRβs likely represented T cell clones that had expanded. We then calculated Spearman’s and Pearson’s correlation coefficients for all high-abundance TCRβ pairs, filling in missing data with the median abundance of TCRβs from each sample. We used median abundance—instead of a pseudocount of 1 or half the minimum abundance detected— because the immense diversity of the TCRβ repertoire means that most detected TCRβs are likely similarly abundant as TCRβs that were not detected. We identified pairs of TCRβs that had high (>0.95) correlation. To identify cohorts of TCRβs that co-correlated, we represented TCRβs as nodes in a network, where nodes were connected by edges if the corresponding TCRβs were highly correlated. We then searched for the maximal network clique (a set of nodes where each node has an edge to all other nodes) using NetworkX. We visually inspected these TCRβ cohorts for evidence of sequencing error, which might have resulted in a high-abundance TCRβ that closely correlated with many low-abundance TCRβs with similar sequences (**Table S2**). To test the significance of TCRβ cohort size, we performed the same analysis on 1000 shuffled datasets. Each shuffled dataset randomly permuted sample labels (i.e., the sampling date) for each TCRβ within each individual.

To test the significance of persistent TCRβ enrichment in (a) public receptors (**Figure 3**) and (b) TCRβs that occurred with many similar receptors (**Figure S13**), we analyzed 10,000 shuffled datasets. For these permutations, we randomly permuted the number of time points at which each TCRβ was observed and repeated the analysis.

We estimated the probability of generation of each TCRβ prior to immune selection using IGoR version 1.1.0 with the provided model parameters for the human TCRβ locus (33).

## DECLARATIONS

### Acknowledgments

We thank Ellen W. Chu for substantive and editorial comments on the manuscript.

### Ethics approval and consent to participate

All procedures were conducted under WIRB protocol ADAP-002 (“Immunology studies of normal healthy individuals”). Subject enrollment and study procedures were directed by Adaptive Biotechnologies

### Consent for publication

Not applicable

### Availability of data and materials

All data are freely available on the immuneACCESS portal of Adaptive Biotechnologies (https://clients.adaptivebiotech.com/immuneaccess).

### Authors’ contributions

ROE, AMS, and HSR conceptualized the experiment and generated the data. NDC and HSB analyzed the data. NDC made figures, and HSB wrote the first draft of the manuscript with input from NDC. All authors contributed to manuscript development.

---

**Figure S1.**
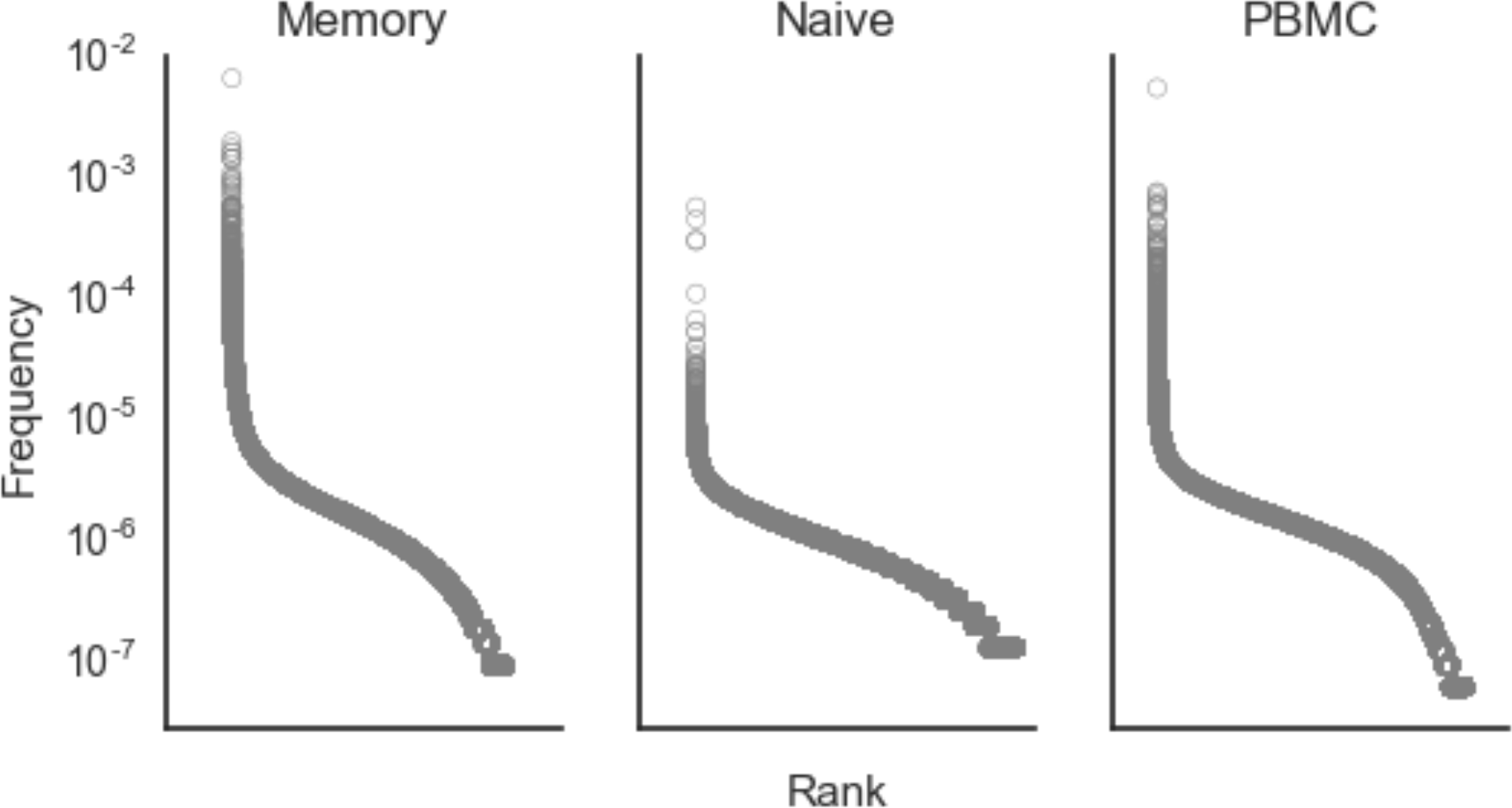
Representative frequency rank plots for memory T cells, naive T cells, and all T cells from PBMCs from Individual 01. As expected, naive T cells had fewer abundant clones than PBMC or memory T cells. In all cases, the majority of TCRβs had abundances around 10^−6^.

**Figure S2.**
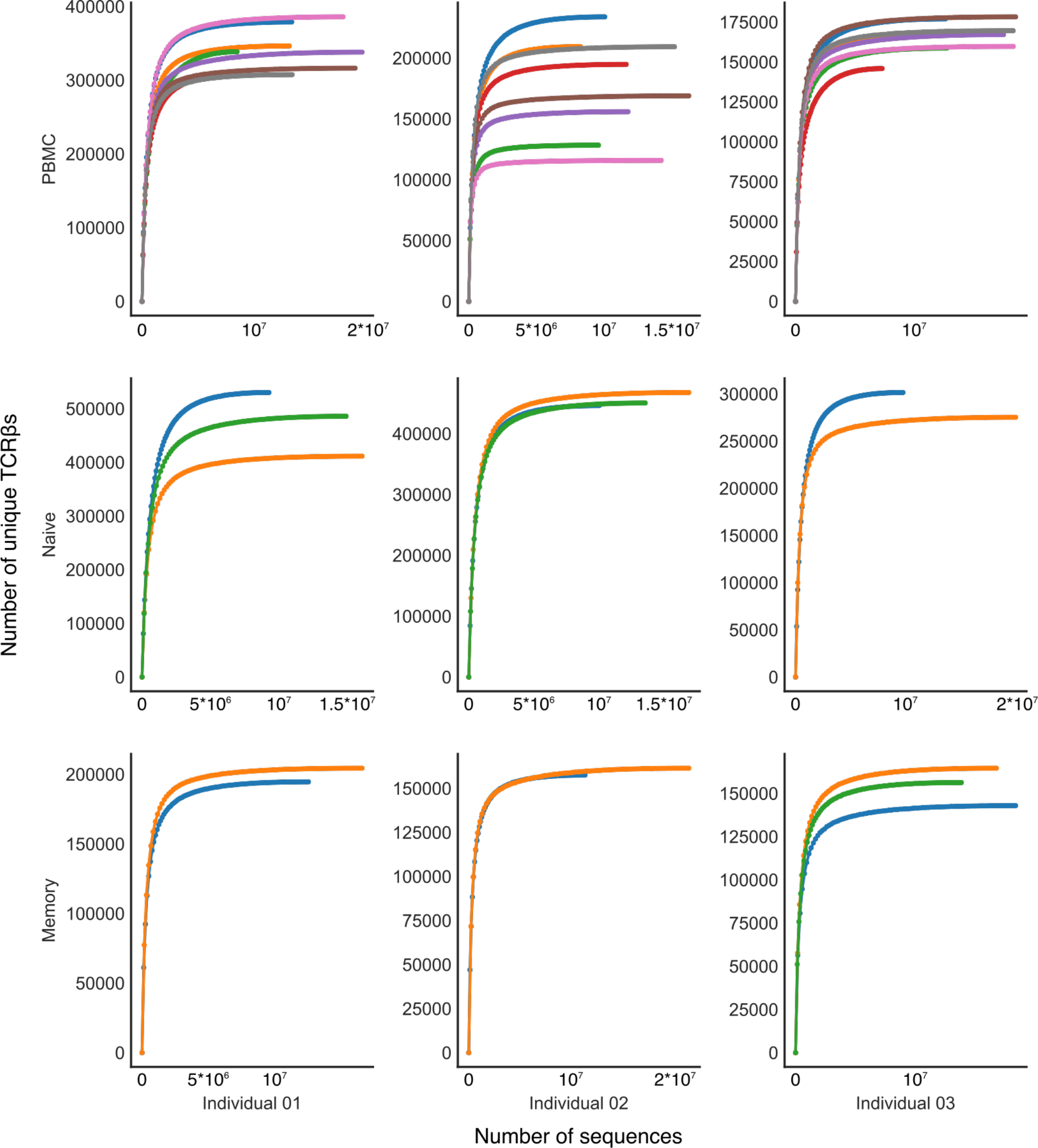
Rarefaction curves for each subject indicate that sample libraries were sequenced well past saturation.

**Figure S3.**
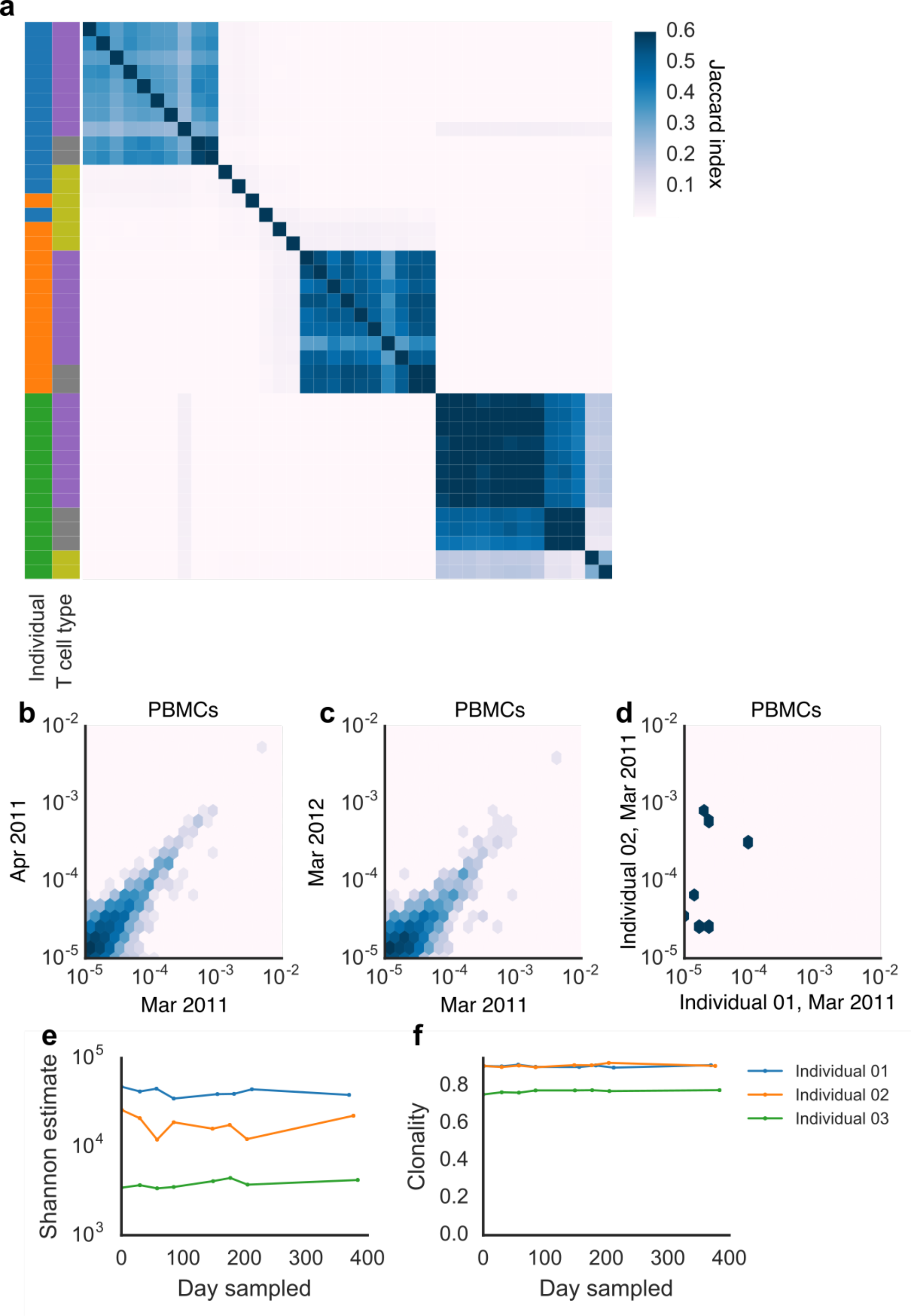
Analyses examining only high-abundance TCRβs agree with results from full-repertoire analysis, suggesting that undersampling likely did not confound our results. (a) A heatmap of Jaccard indexes shows similar clustering of PBMC and memory T cell samples by individual and less clustering of naive T cell samples. Abundances of high-abundance TCRβs in PBMC samples correlated within an individual (individual 01) across time points, including across a month (b, shared TCRβs = 2057, Spearman rho = 0.66902, p < 10^−6^) and a year (c, shared TCRβs = 1390, Spearman rho = 0.59251, p < 10^−6^). High-abundance TCRβs did not appear to correlate across individuals, largely because of lack of shared TCRβs (d, shared TCRβs = 7, Spearman rho = 0.14286, p = 0.75995). Shannon alpha diversity estimate (e) and clonality (defined as 1 – Pielou’s evenness, f) of the TCRβ repertoire were consistent over time.

**Figure S4.**
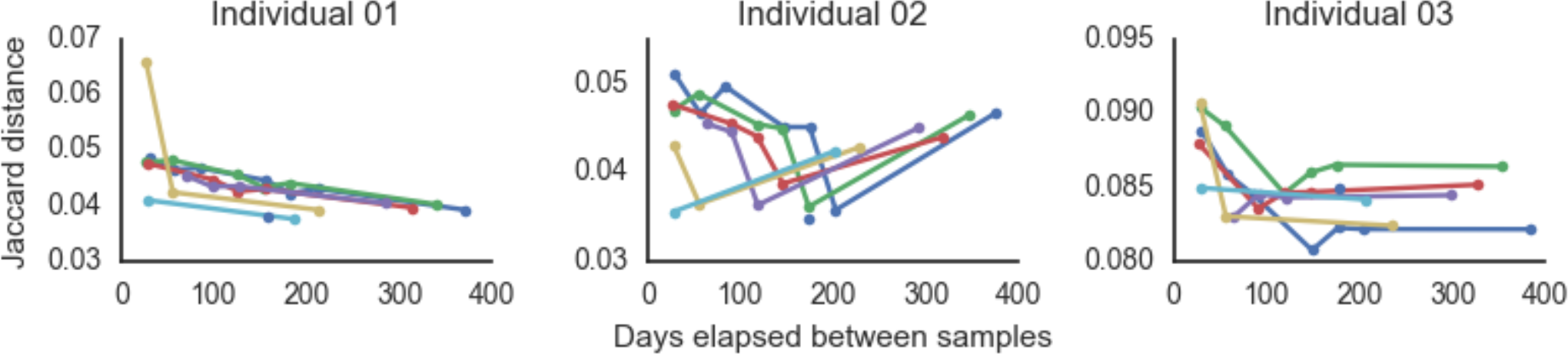
TCRβ repertoire overlap (Jaccard index) often decreases with increasing time between samples, except in Individual 02, where the final time point at one year past the first sample shared more TCRβs with the previous samples.

**Figure S5.**
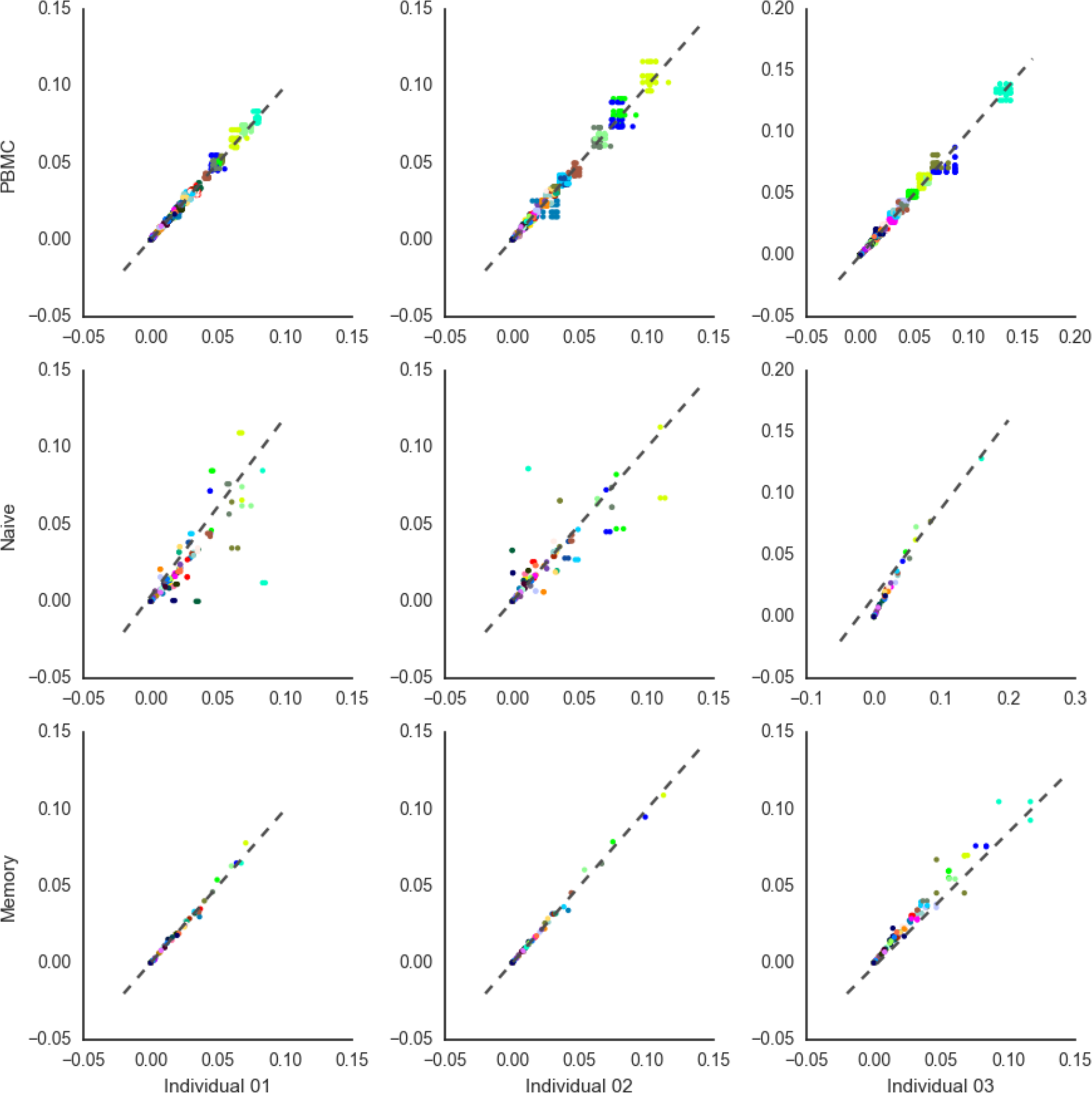
V gene usage across time and cell compartment in all three individuals. Each of nine plots represents each cell type in each individual. Within each plot, different colors represent different V genes, and each dot represents a comparison of the abundances of that V gene from one sample to another. Points that fall near a 1:1 ratio (indicated by the dotted line) are nearly identical in abundance between the two samples considered. These plots indicate that VJ gene usage was generally the same across time points, particularly in total and memory T cells. In naive cells, VJ gene usage varied more.

**Figure S6.**
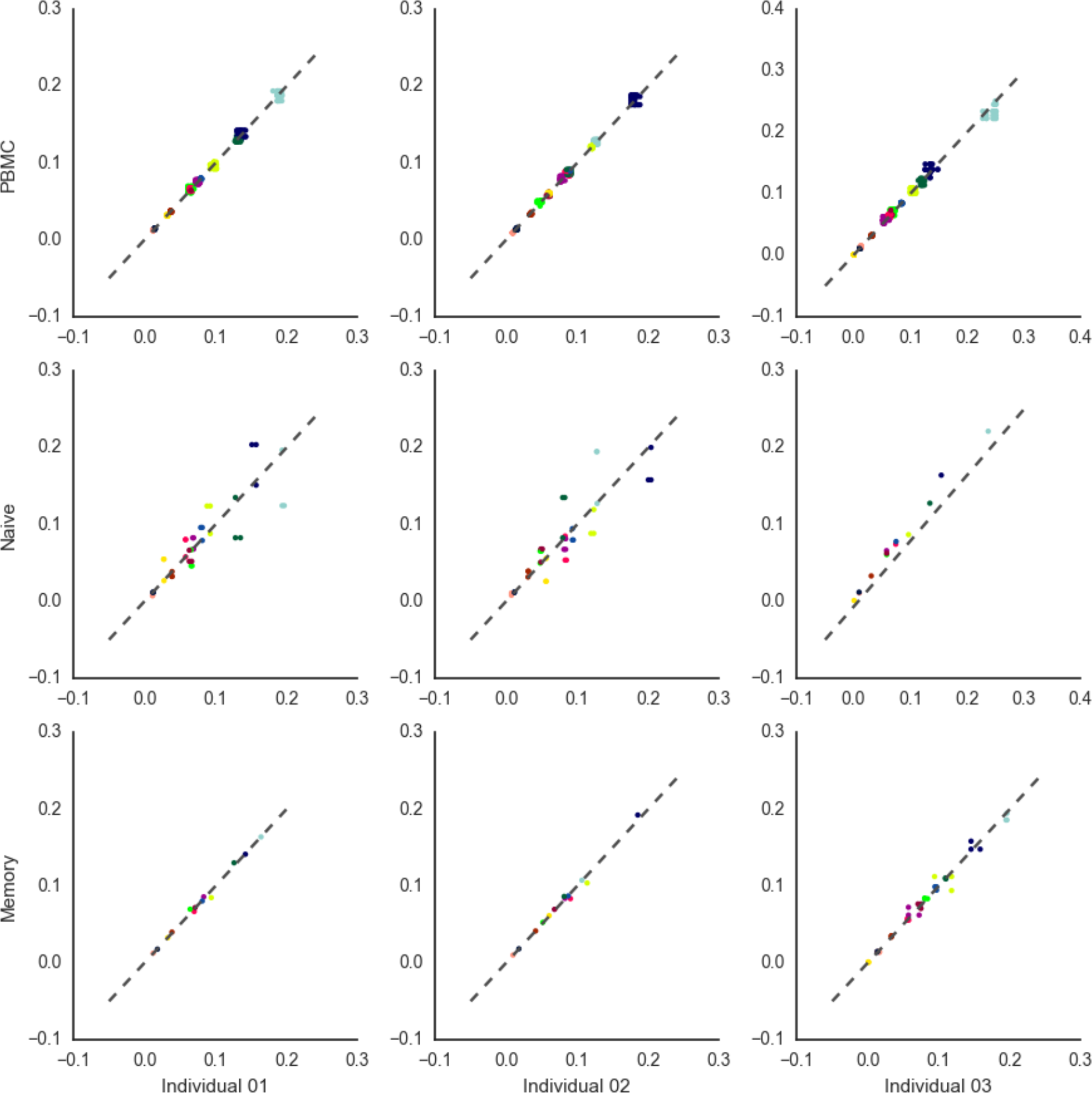
J gene usage across time and cell compartment in all three individuals. Plots are as in **Figure S5**, with similar findings.

**Figure S7.**
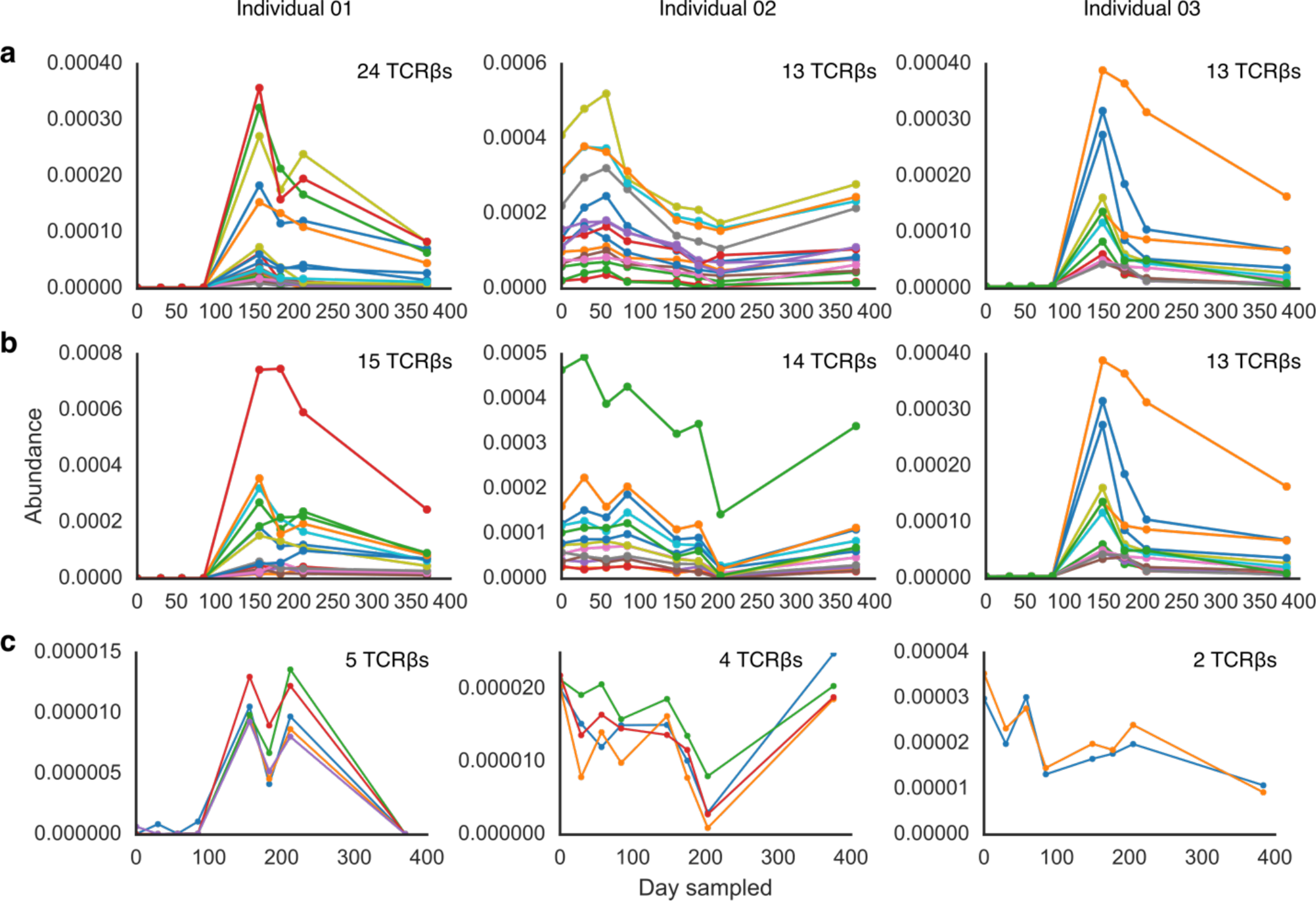
Cohorts of TCRβs exhibit correlated dynamics over time. We found large cohorts of correlating TCRβs by Spearman (**a**) and Pearson (**b**) correlation. Although these TCRβs spanned a range of abundances, we did not observe any clear signs of correlation caused by sequencing or library preparation errors (**Table S2**). We also found smaller cohorts (**c**) of TCRβs with nearly identical abundances whose dynamics also correlated through time. The number of TCRβs found in all cohorts was significant (*p* < 0.001) in a random permutation test (see Methods). These TCRβ cohorts might be an artifact of sampling noise, or they may represent receptors involved in the same immune response.

**Figure S8.**
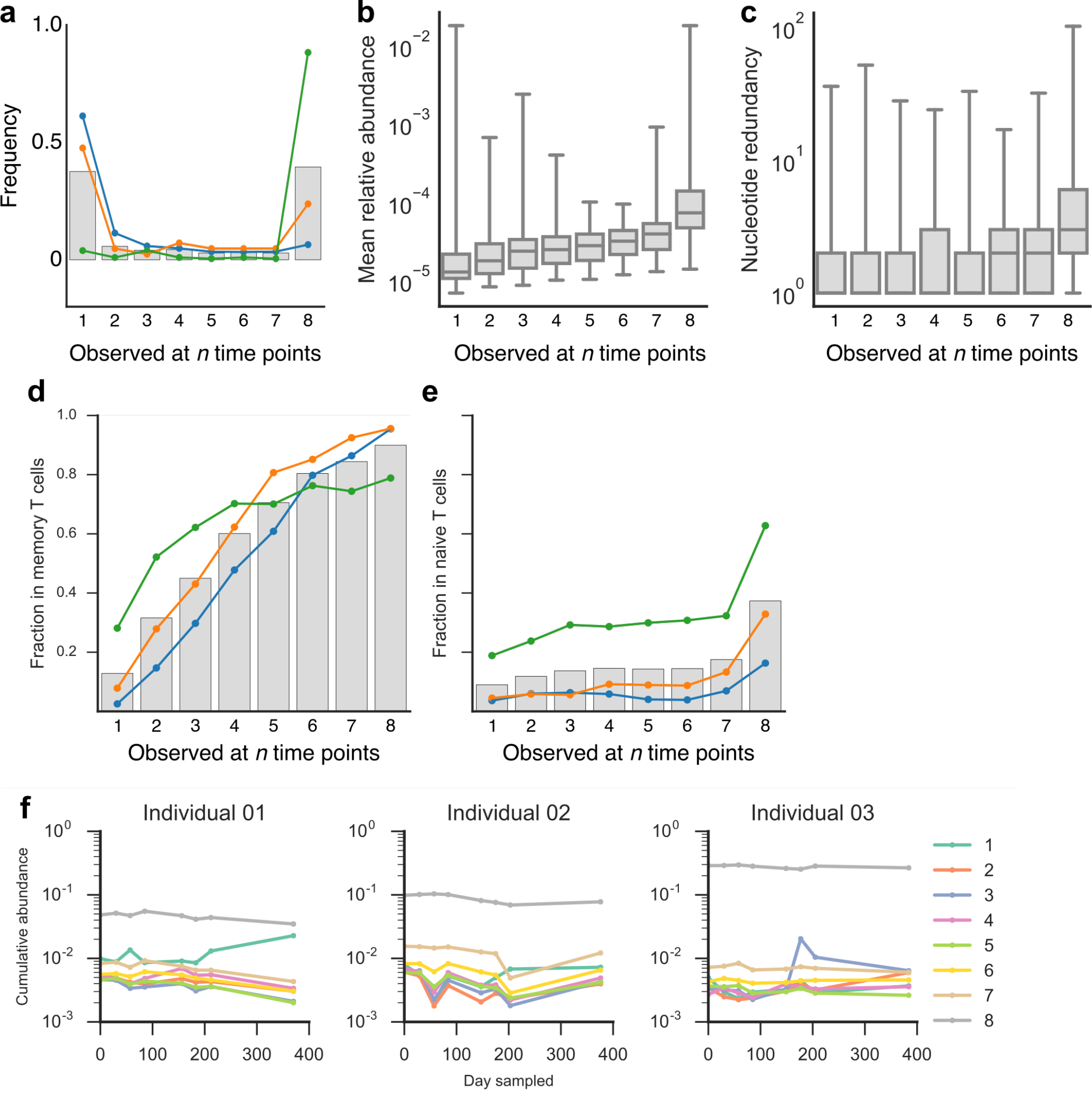
Persistent high-abundance TCRβs exhibit similar patterns as overall persistent TCRβs. (**a**) High-abundance TCRβs had a greater prevalence of persistent TCRβs, although the exact values varied across individuals. Persistent high-abundance TCRβs also showed greater mean abundance (**b**) and nucleotide redundancy (**c**). Persistent high-abundance TCRβs also had higher proportions of TCRβs in common with memory (**d**) and naive (**e**) T cell populations and constituted a stable and significant fraction of overall TCRβ abundance across time (**f**).

**Figure S9.**
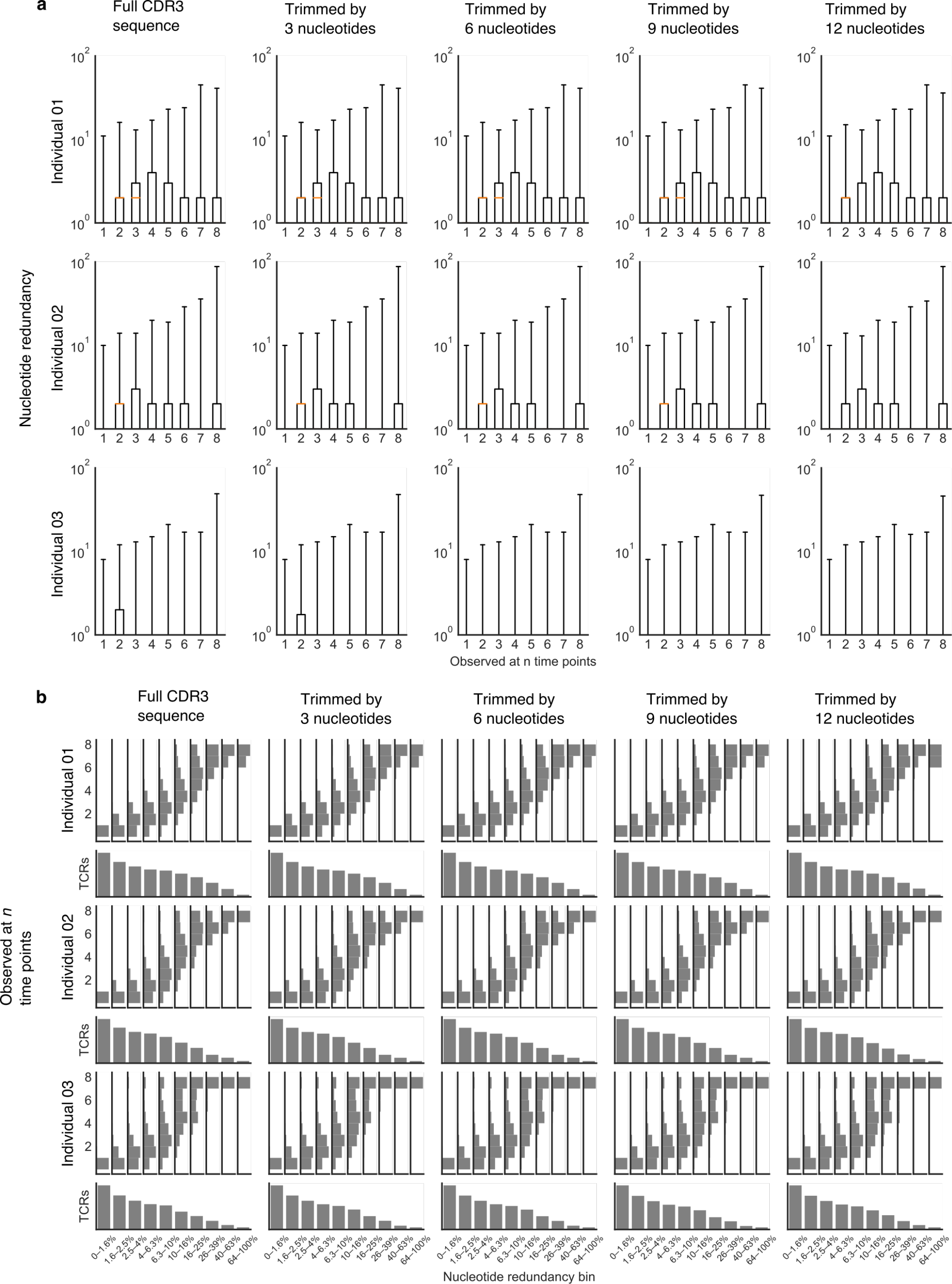
Nucleotide redundancy across individuals and with more stringent assignment of CDR3 sequence (figure supplements Figure 2c). (**a**) Each plot represents nucleotide redundancy for TCRβs that were observed in *n* samples. Rows represent plots for each individual. The leftmost column of plots comprises data from full CDR3 nucleotide sequences as identified by IMGT (as in Figure 2c): we observed that the pattern of increasing nucleotide redundancy in persistent TCRβs was not consistent across individuals. Each of the following columns plot data from CDR3 nucleotide sequences that were progressively trimmed on each end by 3, 6, 9, and 12 nucleotides. We trimmed these sequences because CDR3 sequences identified by IMGT generally capture a number of amino acids—usually one to four at each end of the sequence— that are derived from V and J genes. Nucleotide mutations in these leading and trailing ends are thus less likely to be of biological origin and more likely to be from sequencing error, since we do not expect nucleotides from the V or J genes to be altered during TCR recombination (except for deletions). From these plots, we can observe that nucleotide redundancy is generally stable over different lengths of trimming, suggesting that our data are not skewed by these potential sequencing errors. (**b**) To further examine the relationship between persistence and nucleotide redundancy, we grouped TCRβs into 10 bins according to nucleotide redundancy. Because nucleotide redundancy is extremely skewed—the vast majority of TCRβs are encoded by a single clonotype—we created these bins on a logarithmic scale: the first bin includes TCRβs with nucleotide redundancy values up to 1.6% of the maximum value for each individual; the second between 1.6% and 2.5% of the maximum value; and up to the 10th bin, which includes TCRβs with nucleotide redundancy values between 64% and 100% of the maximum value. For each of these TCRβ bins, we then plotted a histogram of the frequency of TCRβs that were observed at *n* time points. We observe a clear pattern across individuals and trimming lengths: TCRβs with greater nucleotide redundancy tend to occur at more time points, and the most redundant TCRβs are exclusively persistent receptors.

**Figure S10.**
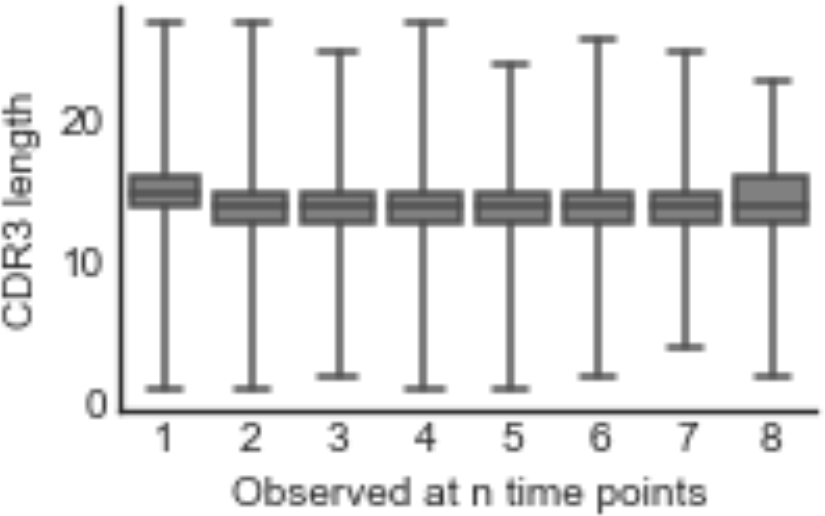
The persistent TCRβ repertoire exhibited little alteration of CDR3 lengths.

**Figure S11.**
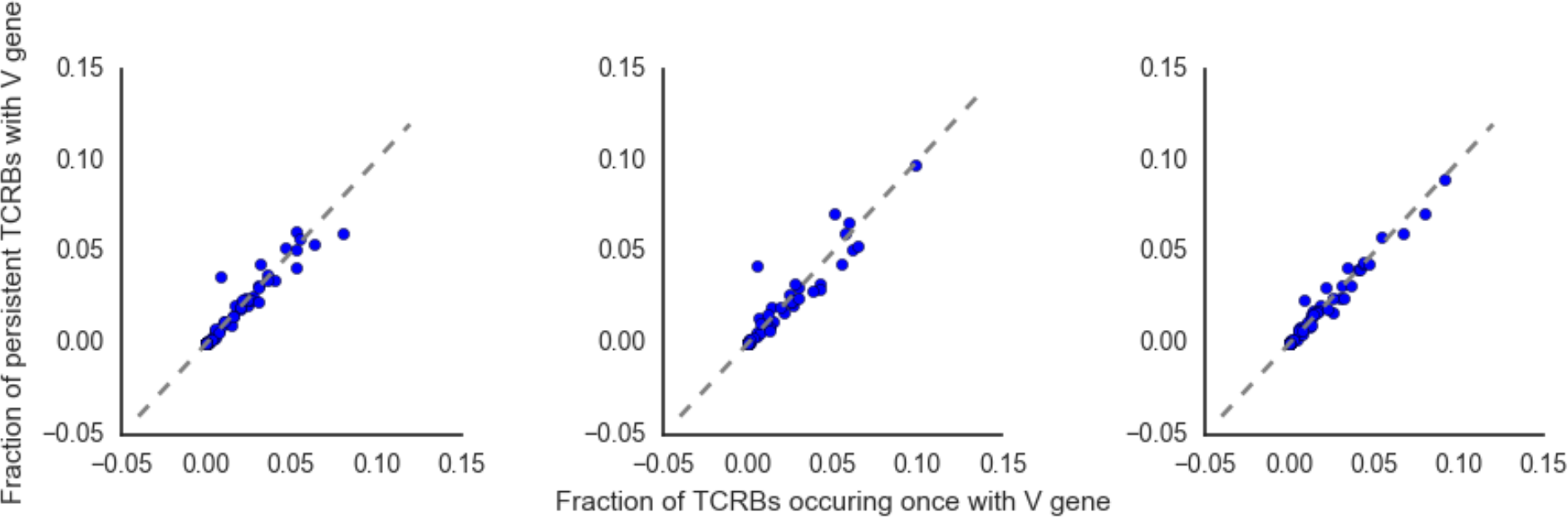
The persistent TCRβ repertoire does not exhibit altered V gene usage. These plots show V gene usage in TCRβs that occurred only once (x-axis) versus in persistent TCRβs (y axis). Each data point represents a single V gene. These values were closely correlated.

**Figure S12.**
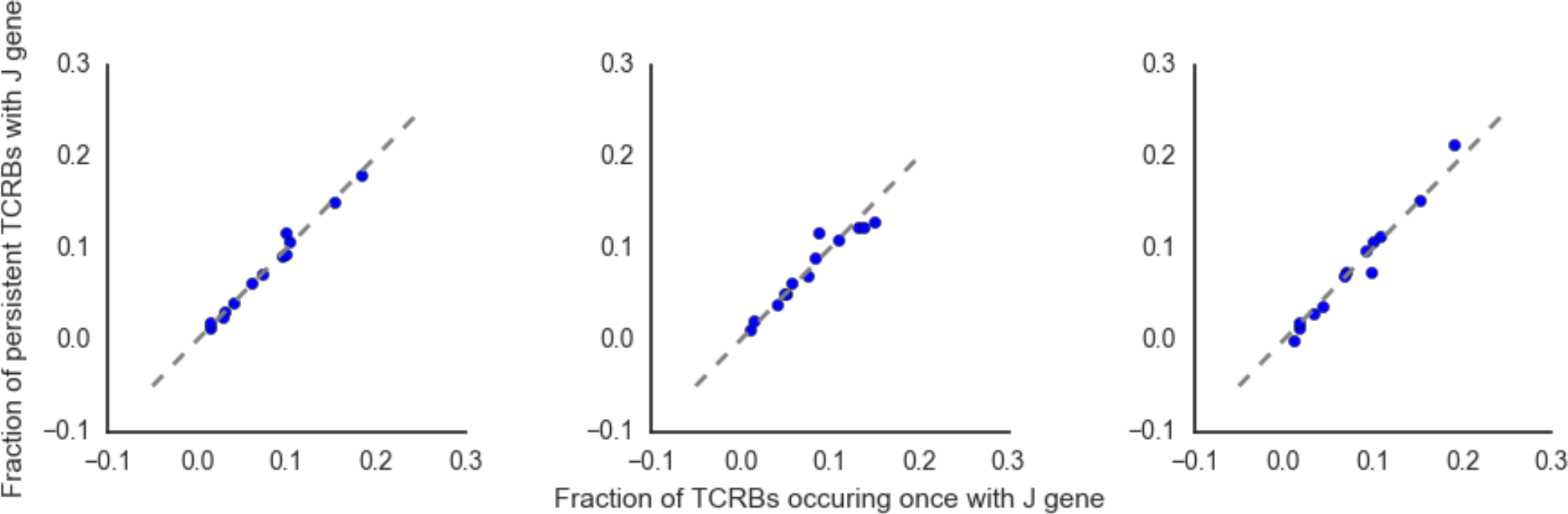
The persistent TCRβ repertoire does not exhibit altered J gene usage. Similar plots as in **Figure S10** indicate that J gene usage is not greatly changed in persistent TCRβs.

**Figure S13.**
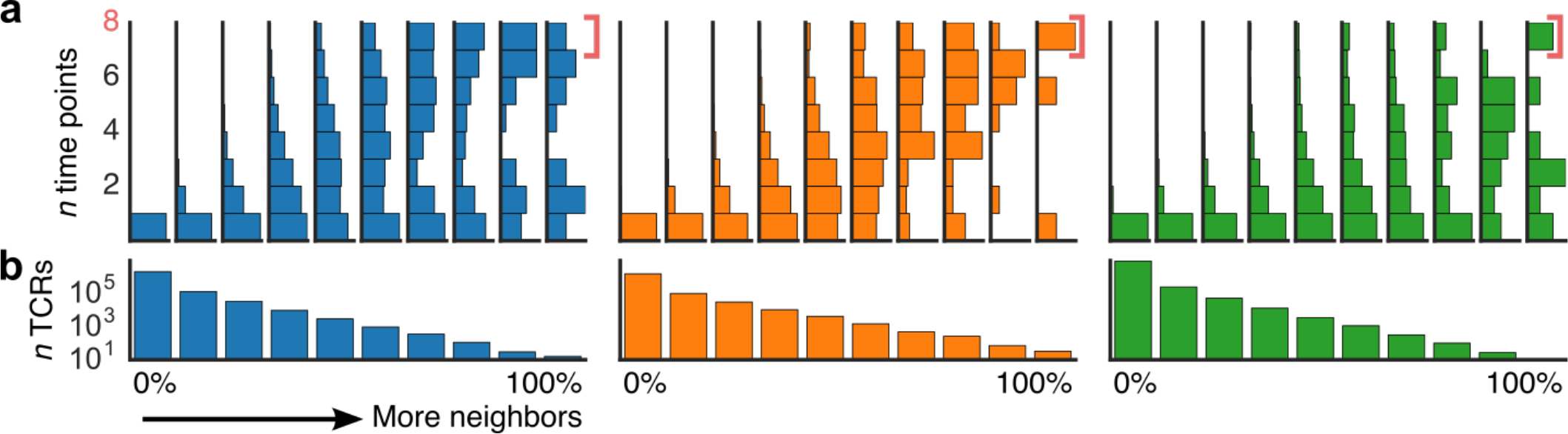
Persistent TCRβs were more functionally redundant. We created a network graph of TCRβs from each individual, drawing edges between TCRβs on the basis of sequence similarity (Levenshtein distances), which reflects antigen specificity. We then grouped TCRβs into decile bins based on the number of neighbors (similar TCRβs) of each TCRβs. For each decile bin, we then evaluated in how many samples each TCRβ occurred in from our time series data. (**a**) TCRβs with higher numbers of neighbors—and thus higher numbers of similar TCRβs observed—tended to have a higher proportion of persistent TCRβs than nodes with few neighbors. (**b**) The number of TCRβs in each neighbor bin. Plot similar to **Figure 3**.

**Figure S14.**
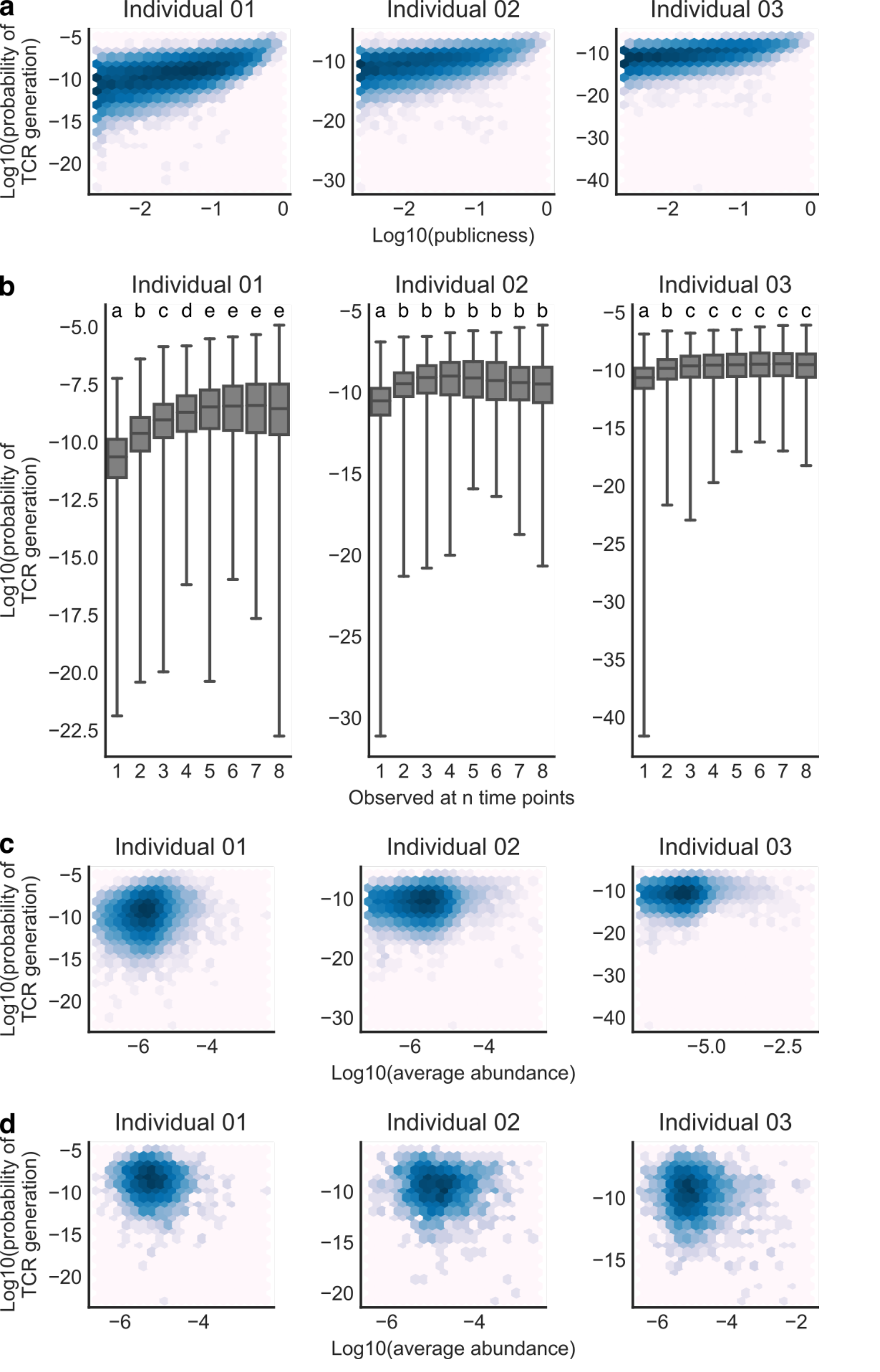
Persistent and public receptors may result in part from TCR recombination biases. (a) As in previous studies, the probability that a given TCRβ was generated correlated closely with publicness in a cohort of 778 individuals. For each individual, only TCRβs occurring in both that individual and the cohort were considered. (number of TCRβs evaluated in individual 01 = 86380, Spearman *rho* = 0.62488, *p* < 10^−6^; number of TCRβs evaluated in individual 02 = 338617, Spearman *rho* = 0.52231,*p* < 10^−6^; number of TCRβs evaluated in individual 03 = 284990, Spearman *rho* = 0.51129, *p* < 10^−6^). (b) TCRβs occurring at more time points tended to have higher generation probabilities, although overall ranges were broad and overlapping. Letters indicate significant differences from all other groups by a Mann-Whitney *U* test (*p* < 0.001). Mean abundance of all TCRβs (c) and only persistent TCRβs (d) did not correlate with generation probability. (All TCRβs: Spearman *rho* = 0.17143, *p* < 10^−6^; Spearman *rho* = 0.05300, *p* < 10^−6^; Spearman *rho* = 0.08208, *p* < 10^−6^). (Persistent TCRβs: number of TCRβs evaluated in individual 01 = 3448, Spearman *rho* = −0.06793, *p* = 0.00007; number of TCRβs evaluated in individual 02 = 1978, Spearman *rho* = −0.04341, *p* = 0.0537; number of TCRβs evaluated in individual 03 = 2965, Spearman *rho* = 0.04552, *p* = 0.01318)

**Table S1.** Overall TCRβ-sequencing statistics per sample: sequencing depth, productive TCRβ sequencing depth, fraction of productive TCRβ sequences, unique V genes identified, unique J genes identified, unique CDR3 sequences, unique TCRβs, unique TCRβ nucleotide sequences.

**Table S2.** Sequence and abundance information for the largest cohort of closely correlated TCRβs identified in each individual by Spearman’s or Pearson’s correlation.

